# Mental development is associated with cortical connectivity of the ventral and nonspecific thalamus of preterm newborns

**DOI:** 10.1101/2020.05.05.078196

**Authors:** András Jakab, Giancarlo Natalucci, Brigitte Koller, Ruth Tuura, Christoph Rüegger, Cornelia Hagmann

## Abstract

The thalamus is a key hub for regulating cortical connectivity. Dysmaturation of thalamocortical networks that accompany white matter injury have been hypothesized as neuroanatomical correlate of late life neurocognitive impairment following preterm birth. Our objective was to find a link between thalamocortical connectivity measures at term equivalent age and two year neurodevelopmental outcome in preterm infants. Diffusion tensor MRI of 58 infants (postmenstrual age at birth, mean (SD), 29.71 (1.47) weeks) was used to trace connections between the cortex and thalami. We found strong correlation between mental developmental index and two complementary measures of thalamocortical networks: connectivity strength projected to a cortical skeleton and pathway length emerging from thalamic voxels (partial correlation, R=0.552 and R=0.535, respectively, threshold-free cluster enhancement, corrected p-value<0.05), while psychomotor development was not associated with thalamocortical connectivity. Post hoc stepwise linear regression analysis revealed that parental socioeconomic scale, postmenstrual age and the duration of mechanical ventilation at the intensive care unit contribute to the variability of outcome. Our findings independently validated previous observations in preterm infants, providing additional evidence injury or dysmaturation of tracts emerging from ventral specific and various non-specific thalamus projecting to late-maturing cortical regions are predictive of mental, but not psychomotor developmental outcomes.

## Background

Preterm birth is a risk factor for a variety of neurodevelopmental impairments. Many children face impairments across multiple neurodevelopmental domains such as cognition, and language, and motor skills [L. J. Woodward et al., 2009]. Prominent problems are seen in executive functions, language development, and behavior [Van’t Hooft et al., 2015] persisting into adolescence [Wehrle et al., 2016] and adulthood [Nosarti et al., 2009]. More recently, a preterm behavioral phenotype has been described that exhibits inattention, anxiety, and social-communication deficits [Johnson and N. Marlow, 2011]. Early brain alterations associated with preterm birth have been described as a complex amalgam of destructive and developmental disturbances and may lead to atypical brain development and ultimately to such impairments. Indeed, there is convincing evidence that very preterm infants are at substantial risk of brain injury in the perinatal period [Volpe, 2009a]. The periventricular white matter (WM) seems to suffer the primary injury, and this WM injury is frequently accompanied by neuronal and axonal disease affecting the cerebral WM, thalamus, basal ganglia, cerebral cortex, brain stem, and cerebellum [Volpe, 2009a; Volpe, 2009b].

The thalamus acts as a key hub for cortical networks and thalamocortical connections, and it is commonly affected, especially in preterm infants, by WM injury, either directly or through maturational disturbance [Boardman et al., 2006; Nosarti et al., 2008]. The development of thalamocortical connections during the mid- and late gestational periods plays a critical role in shaping brain connectivity during prenatal and early postnatal life [Ghosh et al., 1990; Kostovic and Judas, 2010; McQuillen and D. M. Ferriero, 2005]. The ingrowth of thalamocortical connectivity is a crucial milestone for the sensory-expectant organization and functional specialization of the cerebral cortex [Kostovic and M. Judas, 2002; Molliver et al., 1973; Molnar et al., 1998]. The topographical organization of thalamic connections is established early after side branches reach the cortex radially from the subplate [Catalano et al., 1996; Ghosh and C. J. Shatz, 1992], and their organization is further shaped by activity-dependent synaptic interaction [Molnar et al., 2002]. Thalamocortical fibers begin to relocate to the cortical plate around the 24^th^ week of gestation in sensory and later in association cortices [Kostovic and M. Judas, 2006; Kostovic and M. Judas, 2007]. Crucially, this is the period of biological vulnerability due to prematurity. Together, these developments provide strong biological evidence that the maturation of thalamocortical connectivity is affected by premature birth and that injury to the thalamocortical circuitry may indirectly affect cortical functional specialization.

Evidence is emerging for alterations in thalamocortical connectivity after preterm birth at term-equivalent age [Ball, Boardman et al., 2013; Fischi-Gomez et al., 2016a; Kelly et al., 2016; Pandit et al., 2014; van den Heuvel et al., 2015] and that such alterations are associated with later neurodevelopment [Ball et al., 2015; Fischi-Gomez et al., 2016b]. Hence, both injury to the thalamus and alterations to the thalamocortical connections might impact the cognitive abilities of preterm infants. For example, the role of the mediodorsal thalamus in distinct cognitive behavior that relies on various prefrontal regions has been described in patients with schizophrenia [N. D. Woodward et al., 2012], and anatomical variability of the connections of the mediodorsal nucleus has been linked to the variation of executive functions in adults [Jakab et al., 2012].

A better understanding of the pathomechanism of injury to the thalamocortical circuitry after preterm birth is essential to elucidate how its alteration may contribute to cognitive impairment. The present study was designed to provide further evidence and independent validation for the theory that the dysmaturation of thalamocortical connectivity in preterm infants is predictive of later neurodevelopmental outcomes. We hypothesize that the neurodevelopmental sequelae mirror the emergence of thalamocortical connectivity topography and that the thalamocortical connectivity of late-maturing regions is therefore predictive of cognitive outcomes. Testing this hypothesis required us to identify the sub-thalamic anatomical locations with the strongest putative correlation between thalamocortical connectivity and later life cognition.

## Methods

The preterm infants in this study represent a subgroup of infants enrolled in a randomized, double-blind placebo-controlled, prospective multicenter study titled “Does erythropoietin improve outcome in preterm infants?” (NCT00413946) that were examined by means of a cranial MR at term equivalent age. This subgroup of infants has been described previously [Jakab et al., 2019; O’Gorman et al., 2015]: the main criterion for subjects to be enrolled in this subgroup was the availability of good quality cerebral diffusion tensor imaging (DTI) data. Fifty-eight preterm infants with mean (SD) gestational age at birth of 29.75 (1.44) weeks and at scanning of 41.09 (2.09) weeks) were included in this analysis. The characteristics of the infants are described in Table 1. Socioeconomic status was estimated by a validated 12-point socioeconomic score based on maternal education and paternal occupation, and the infants were classified into higher class (score 2-5), middle class (6-8), and lower class (9-12) ([Largo et al., 1989]. The local ethical committee approved the project (KEK StV-36/04), and the Swiss drug surveillance unit (Swissmedic, 2005DR3179) approved the study. The trial was registered at ClinicalTrials.gov (number NCT00413946). The data that support the findings of this study are available on request from the corresponding author. The data are not publicly available due to privacy or ethical restrictions.

**Table 1.** Study population.

| Characteristics | Value |
| --- | --- |
| Mean (SD) GA at births (weeks) | 29.71 (1.47) |
| Mean (SD) birthweight (grams) |  |
| Mean (SD) age at MRI | 41.09 (2.1) |
| Female, n (%) | 39 (67%) |
| Chronic lung disease, n (%) | 9 (15.5%) |
| ROP† >3 | 3 (5.1%) |
| Necrotizing enterocolitis, n (%) | 1 (5.2%) |
| Median (range) SES | 5 (1-10) |

† retinopathy of prematurity

### Outcome assessments

Two year neurodevelopmental outcomes of the whole study population have previously been published [Natalucci et al., 2016]. A developmental assessment using the Bayley Scales of Infant Development, second edition (BSID-II, [Bayley, 1993]) was performed at a mean age of 23.4 (2.33) months by experienced developmental specialists. Two outcome measures, were calculated: mental development index (MDI) and psychomotor development index (PDI). The developmental specialists were blinded to the MRI findings.

### MRI acquisition

Neonatal cerebral MRI was performed at term-equivalent age with a 3.0 T GE scanner (GE Medical Systems), using an eight-channel receive-only head coil All infants were scanned during natural sleep using a vacuum mattress. Ear plugs and minimuffs were applied for noise protection. During the scanning, oxygen saturation was monitored, and a neonatologist and a neonatal nurse were present.

Diffusion tensor imaging (DTI) was acquired using a pulsed gradient spin echo planar imaging sequence with TE/TR: 77/9000 ms, field of view = 18 cm, matrix = 128 × 128, slice thickness = 3 mm. For each infant, 21 non-collinear gradient encoding directions with b = 1000 and four interleaved b = 0 images were acquired. The DTI data (n=58) used in our study is part of a previously reported dataset (n=140) [Natalucci et al., 2016].

### Image post processing

DTI data were visually controlled for artifacts. Image frames and the corresponding entries in the b-matrix and b-value descriptor files were removed from further analysis if head movement of the infant caused extensive signal dropout throughout the brain in the given frame in more than one slice along the superoinferior axis. We discarded these data if the newborn woke up or moved excessively during the DTI scan. The entire dataset was excluded if more than three diffusion-weighting gradient volumes were corrupted by motion artifacts. We experienced high drop-out rate because DTI was acquired towards the end of the examination, and many infants woke up or moved excessively (excluded data sets based on subjective assessment: n=78). Four infants were excluded because of cystic lesions (n=4). In the remaining cases, the number of removed image frames (due to excessive patient motion) was recorded as a confounder.

We used a custom script written in Bash language for Linux to process the neonatal DTI images. During the first step of post processing, spurious image shifts originating from eddy currents were corrected with the *eddy* command in the Functional Magnetic Resonance Imaging of the Brain Software Library (FSL) software (volume-to-volume reconstruction). Afterwards, Rician noise filtering of DTI was performed using the command line module *JointRicianLMMSEImageFilter* in Slicer 3D [Aja-Fernandez et al., 2009]. Diffusion tensors and scalar maps were estimated in each voxel using the *dtifit* program in FSL with weighted least squares estimation.

Next, a standard space fractional anisotropy (FA) template image was created by the three-step coregistration of 40 FA images to the 42-week template of the ALBERTs neonatal atlas [Gousias et al., 2012]. The source of the DTI data was another study and consisted of normally developing newborns imaged at term equivalent age (mean, SD (range) postmenstrual age (PMA) PMA of the subjects: 42.5 ± 1.9 (39 – 48.7) weeks, 20 females and 20 males). First, the S0 (equivalent to T2-weighting) images were aligned to the T2-weighted template from the ALBERTs atlas using a 12-degrees-of-freedom linear registration in FSL. The resulting transformation matrix was used to coarsely align the FA image of each individual to the template, and a mean linear FA template was saved by averaging the resulting images. Then, each normal subjects’ FA image was linearly and then nonlinearly coregistered with the mean linear FA template to achieve a sharper, nonlinear FA template. For the nonlinear registration step, the reg_f3d command was used in the NIFTIREG software package. The deformation grid vertex distance was 9 mm; the regularization criterion was the weight of the bending energy penalty term set to 0.05; and the smoothness of the deformation fields was achieved by a Gaussian smoothing with a spherical kernel of 4 mm diameter. The FA image of each individual was used to guide the nonlinear transformation that achieved overlap between the DTI space and the standard space FA template. This transformation was used to propagate three binary masks from the ALBERTs atlas to the subject space: whole cortex map, whole thalamus map, and cerebrospinal fluid spaces.

### Thalamocortical connectivity mapping

The association between cognitive and thalamocortical development was tested with three complementary analysis approaches. The first analysis was based on Ball et al.’s [Ball et al., 2015] method, the projection of the thalamocortical connectivity strength (TC) to a cortical skeleton. Second, we analyzed the thalamic origins of the fiber pathways of the first step by measuring the apparent length of thalamocortical fiber pathways emerging from every voxel of the thalamus (TC length). Third, we quantified how strongly each thalamic nucleus is connected to the cortical regions in which thalamocortical connectivity was found to be predictive of 2-year cognitive development. As diffusion tractography is unable to differentiate between afferent and efferent connectivity, we use the expression “thalamocortical” for both corticothalamic and thalamocortical connections throughout the manuscript.

First, thalamocortical anatomical connectivity was mapped with probabilistic diffusion tractography in the FSL software (*Probtrackx2* command). First, the possible orientation distribution priors were estimated using a Bayesian method (*bedpostx* script), allowing for two fiber populations per voxel. Probabilistic tractography was seeded from a region of interest covering the supratentorial cortical gray matter (cortex label of the ALBERTs atlas; [Gousias et al., 2012]). From each cortical voxel, 5000 probabilistic tracing particles were emitted and a mask of cerebrospinal fluid spaces was set to terminate any tracing samples reaching the cerebrospinal fluid. The thalamus mask was used as target for tractography; during this step, only those pathways were kept that passed through the thalamus. A counter in each cortical voxel increased whenever the tracing particle from that voxel reached the thalamus, resulting in seed-to-target probability maps. Secondly, whole-brain maps depicting the connectivity strength (*fdt_paths*) and the mean length of probabilistic fiber pathways in each voxel were exported for further analysis.

We used a modified version of the tract-based statistics (TBSS; [Smith et al., 2006]) for the analysis of thalamocortical connectivity across subjects. TBSS offers a way to overcome the limitation of compromised cross-subject registration during group analysis by providing an alignment-invariant representation of fiber tracts. Instead of the standard method, which consists of generating a representation of the fiber tracts based on fractional anisotropy images, we calculated the centerline of the cortical mantle based on a groupwise cortex probability map of the 42-week template in the ALBERT atlas. The strength of thalamocortical connectivity at each cortical voxel was projected to the nearest cortical centerline voxel using the approach described in the TBSS literature [Smith et al., 2006].

### Anatomical subdivisions of the newborn thalamus

As there are currently no cyto- or myeloarchitecture-based atlases for the human newborn brain, we utilized the statistical shape model version of the adult Morel atlas to localize thalamic subdivisions in the newborn brain [Krauth et al., 2010]. This atlas incorporates histological definitions of nuclei based on seven postmortem samples. We delineated the visible macroscopic borders of the thalamus on the T2-weighted, gestational age-specific template of the ALBERT atlas. First, each training sample was aligned to the template by minimizing the surface distance between the MRI and histology-based thalamus outlines with a thin-plate registration algorithm. The statistical shape model stores anatomical correspondences between the outer borders of the whole thalamus and each thalamic nucleus in each training sample, and this function was used to predict the unobservable geometry (within-thalamus borders) for the rest of the thalamus. Confidence region maps for nuclei were generated by projecting the matching uncertainty of each vertex into a separate grayscale image for each nucleus. A label value in each voxel identified the most probable thalamic nucleus based on the highest confidence (using the *find_the_biggest* command in FSL). The delineation and statistical shape-matching procedure was performed in the NeuroShape extension of the Slicer 3D software, version 4.8.10. The NeuroShape software is based on the method described in our previous work [Jakab, Blanc, Berenyi and Szekely, 2012]. The resulting masks (**Figure 1**) were used to localize the thalamic specificity of our findings.

**Figure 1.**
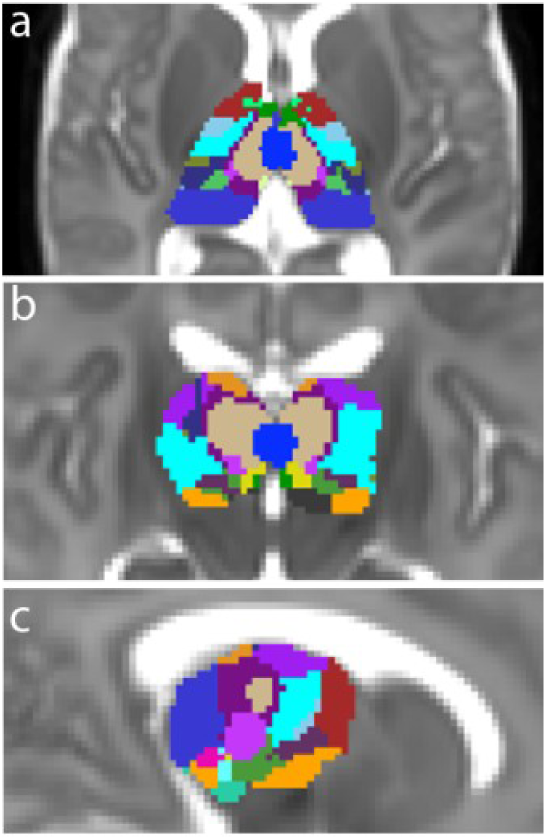
Neonatal thalamus atlas derived from the statistical shape-model-based adaptation of an adult thalamus atlas to a T2-weighted neonatal atlas template corresponding to the 42^nd^ postmenstrual week. (a-c): axial, coronal and sagittal views of the Atlas.

### Statistical analysis

First, we used multivariate linear regression to test whether MDI and PDI correlated with the TC values that were projected onto the cortical skeleton. Statistical analyses were completed using the threshold-free cluster enhancement method (TFCE) [Smith and T. E. Nichols, 2009] with randomized nonparametric permutation testing implemented in the *randomise* program [Winkler et al., 2014], version 2.9, part of the FSL software package, build 509 [Jenkinson et al., 2012]. The default parameter settings of TFCE were used with 5000 permutations. For the initial analysis, familywise error-corrected p≤0.05 was accepted as significant. The same statistical test was then performed for the TC length of all voxels inside the thalamus. Next, post hoc analysis was performed using stepwise linear regression in IBM SPSS V22 (IBM, Armonk, New York) to select any further demographic or clinical parameters that could influence the correlation between TC, TC length, and the cognitive scores. During the stepwise selection of variables from a pool of demographic and neonatal clinical parameters (**Supplementary Table 1**), the probability of F was set to 0.05 for a variable to enter and 0.1 for removal. If clinical variables were selected, we performed the first test again with and without these variables to evaluate whether they are mediators or confounders of the effect.

## Results

### Cognitive scores

The mean (SD) age at the neurodevelopmental assessment was 23.4 (2.33) months. The mean (SD) mental developmental index of the BSID II was 94.2 (14.3), and the mean (SD) psychomotor index was 92.1 (13.3). There was a moderate, negative correlation of SES with MDI (PMCC, - 0.463) but not with PDI (PMCC, 0.0306).

### Correlation between thalamocortical connectivity strength (TC) and outcome at 2 years of age

Our statistical models were corrected for PMA, as diffusion anisotropy and TC might change rapidly during early development, as was also confirmed by our data (**Supplementary Figure 1**). The variable that described the assignment of infants to EPO-treated or placebo groups was not used as a confounder, as neither did EPO treatment predict outcome in this subgroup nor correlated with thalamocortical connectivity in the same cluster-level analysis (TFCE, corrected p>0.05).

First, we found that the 2-year psychomotor development index (PDI) was not correlated with TC in a model adjusted for PMA. The cortically projected TC was significantly correlated (p<0.05) with MDI in predominantly frontal lobe areas, extending from the precentral gyrus to the dorsolateral prefrontal cortex anteriorly and the middle frontal gyrus ventrally (Figure 2/a). Additionally, clusters of correlation were observed in the medial orbitofrontal cortex and the right inferior parietal lobule (**Figure 2/a, c**). To capture the effect size and the contribution of the explanatory variables (PMA and thalamocortical connectivity strength) to the variance of MDI, the average thalamocortical connectivity strength over the voxels which were correlated with MDI was entered into a multivariate linear regression model. We found that TC was strongly, positively correlated with MDI (partial correlation=0.55), while PMA was weakly, negatively correlated with MDI; this second correlation did not reach significance in the model (**Table 2**).

**Table 2.** Correlation between cortically projected thalamocortical connectivity (TC) in the prefrontal cluster and mental development index at two years.

| Predictors† | Standardized Beta | t | p | Zero-order correlation | Partial correlation | Part correlation |
| --- | --- | --- | --- | --- | --- | --- |
| Mean TC‡ | 0.565 | 4.682 | 0.000022 | 0.517 | 0.552 | 0.550 |
| PMA | -0.210 | -1.742 | 0.088 | -0.082 | -0.239 | -0.205 |

† Overall model fit: $F = 11.2$ , adjusted $R^2 = 0.282$ , $P < 0.001$ .
‡ Mean TC calculated from the TFCE-adjusted significant clusters in the cortical skeleton ( $p < 0.05$ )

**Figure 2.**
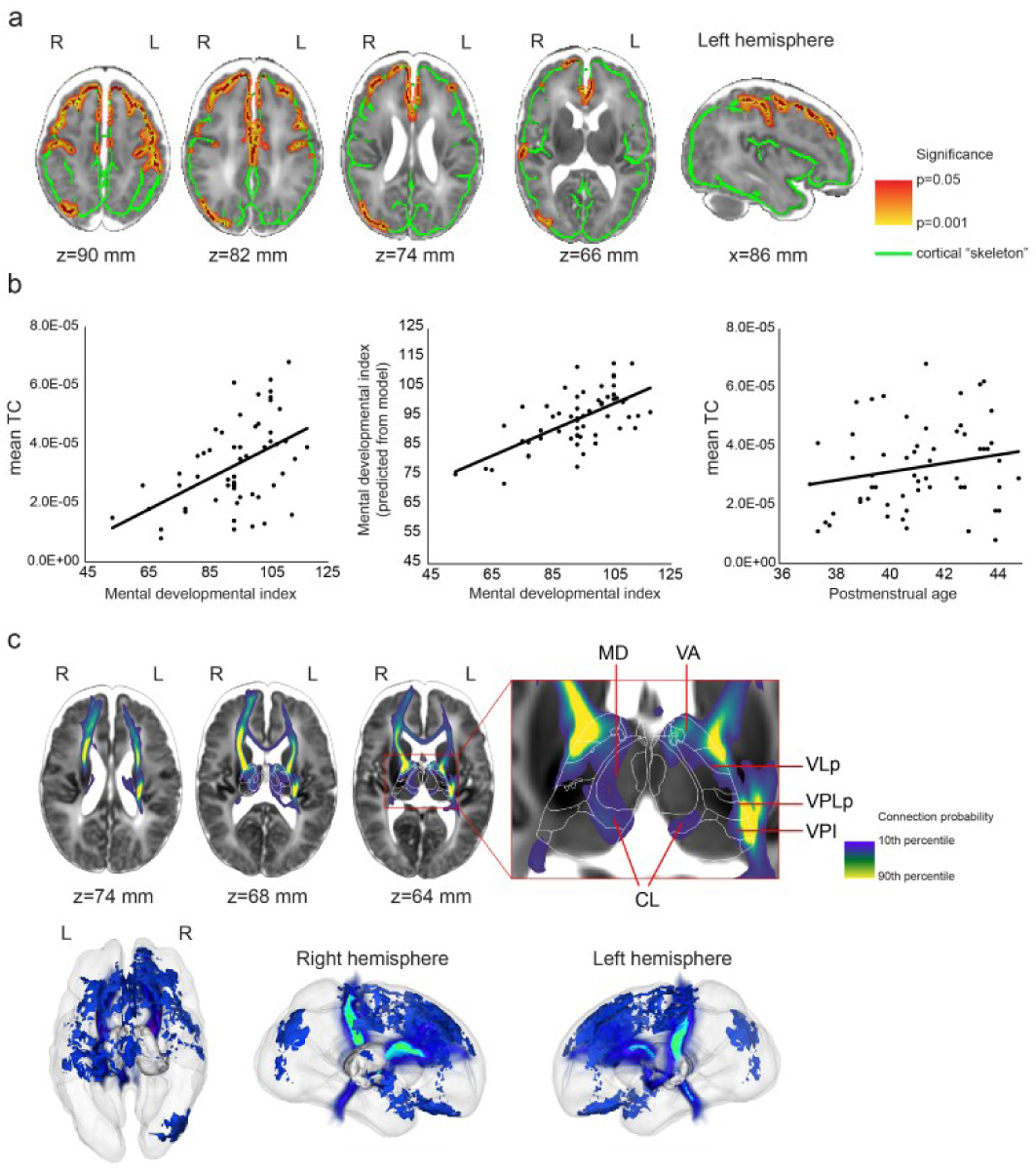
Thalamocortical connections at term equivalent age significantly correlated with 2-year MDI scores in preterm born infants. (a) parts of the cortex where TC and MDI were significantly correlated were marked with red. The significant voxels are dilated to increase visibility, and the dilated voxels are displayed as an overlay colored yellow and red. The cortical skeleton is displayed as green overlay on a T2-weighted MRI template of the 42-week newborn brain. (b) regression plots of the TC–MDI correlation and gestational age dependence of TC averaged over the significant voxels. (c) projection of fiber pathways from the cortical skeleton voxels, where TC was correlated with 2-year mental development. Top images: 2D cross-sectional images where probabilistic, group-average pathways were fused with the MRI template, bottom images: 3D representation of the population mean in a glass brain. Thalamic nuclei abbreviations are given in **Supplementary Table 2**

Next, we used a post hoc, stepwise, linear regression analysis on the mean TC values from the previous experiments to search for possible confounders or mediators of the effect from clinical and demographic variables. In this test, four variables were selected in the model that best explained the variance of MDI: mean TC, days of mechanical ventilation, SES, and PMA. Interestingly, gestational age (GA) at birth was not found to explain the variance of MDI. The association between mean TC and MDI remained strong (partial R=0.608, p<0.001) after adjusting for these variables (**Table 3**). The total time of mechanical ventilation in days was found to be a partial mediator of the effect of mean TC on MDI: after removing this variable from the model, the overall fit was considerably reduced (F=12.91, no adjustment, vs. F=17.01 - adjusted for mechanical ventilation), and the adjusted R^2^ was lower than with the variable in the model (0.407 vs. 0.552). The test statistics of clinical and demographic variables that were either entered or excluded from the final statistical model are given in **Supplementary Table 1**.

**Table 3.** Correlation between cortically projected thalamocortical connectivity (TC) and mental development index at 2 years after adjusting for clinical and demographic variables.

| Predictors† | Standardized Beta | T | p | Zero-order correlation | Partial correlation | Part correlation |
| --- | --- | --- | --- | --- | --- | --- |
| Mean TC‡ | 0.518 | 5.309 | 0.000003 | 0.517 | 0.608 | 0.493 |
| Time of mechanical ventilation | -0.390 | -4.100 | 0.00016 | -0.402 | -0.509 | -0.381 |
| SES | -0.304 | -3.162 | 0.0027 | -0.463 | -0.415 | -0.294 |
| PMA | -0.259 | -2.684 | 0.01 | -0.082 | -0.361 | -0.249 |

† Overall model fit: $F = 17.011$ , adjusted $R^2 = 0.552$ , $P < 0.001$ .
‡ Mean TC calculated from the TFCE-adjusted significant clusters in the cortical skeleton ( $p < 0.05$ )

### Thalamic specificity of probabilistic diffusion tractography results

In each subject, we performed probabilistic diffusion tractography in standard atlas space from the voxels on the cortical skeleton where TC was significantly associated with MDI, and we then analyzed the trajectory and thalamic projection of these connections (**Figure 2/c**). By using the standard-space neonatal thalamus atlas, we quantified the probability of each nucleus being connected to the significant voxels (**Table 4**) by dividing the number of probabilistic streamlines entering all the voxels of a given nucleus by the total number of streamlines. The TC fibers most correlated with MDI were localized in the ventral and anterior nuclei groups: namely, ventral-anterior, ventrolateral, ventromedial, and ventral posteromedial nuclei. Lower connection probability was found for the laterodorsal, anteroventral, and centrolateral nuclei. We observed a moderate left–right asymmetry in the thalamic projections: on the right side, nonmotor or sensory-related nuclei, such as lateral pulvinar or medial geniculate nucleus (MGN), were found to be connected to the significant cortical voxels.

**Table 4.**
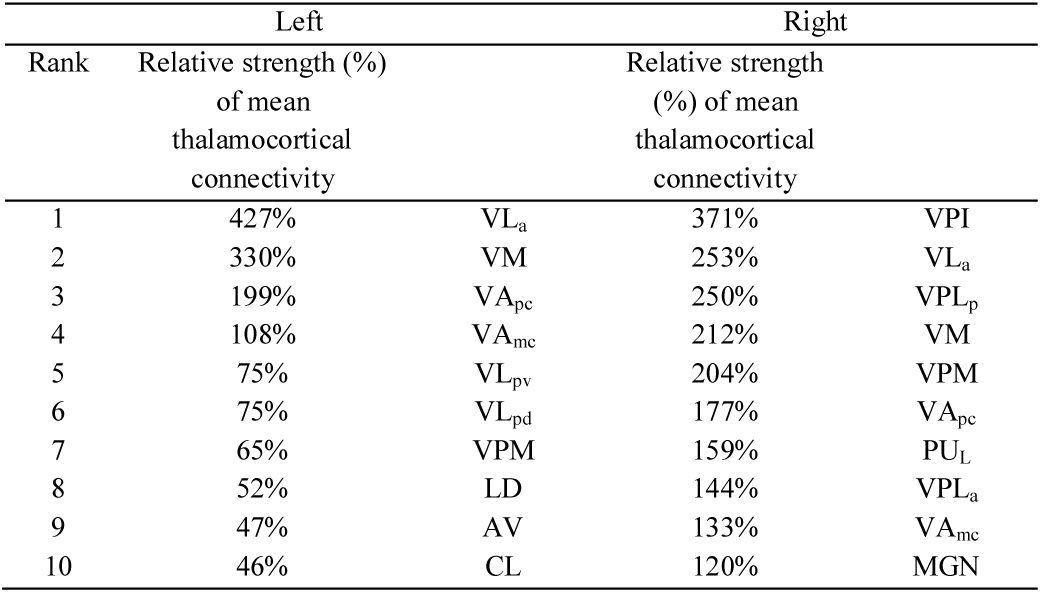
Intrathalamic specificity of our findings: localization of fibers emerging from the cortically projected TC significantly correlated with MDI (details and abbreviations of the thalamic nuclei is given in **Supplementary Table 2**).

### Correlation of thalamocortical fiber length and mental developmental index

We tested whether the length of probabilistically traced thalamocortical fibers (=TC length) was correlated with 2-year cognitive scores. We found that TC length was significantly correlated with (p<0.05) MDI (**Figure 3/a**). The post hoc multivariate linear regression analysis revealed that TC length was strongly correlated with MDI, while PMA was weakly, negatively correlated with MDI (**Table 5**).

**Table 5.**
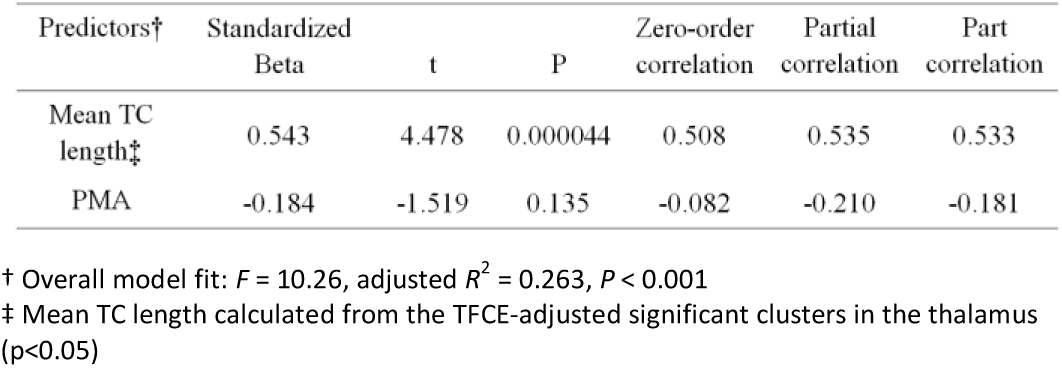
Multivariate linear regression analysis of the correlation between mean thalamocortical fiber length (TC length) in the ventroanterior thalamus and MDI

**Figure 3.**
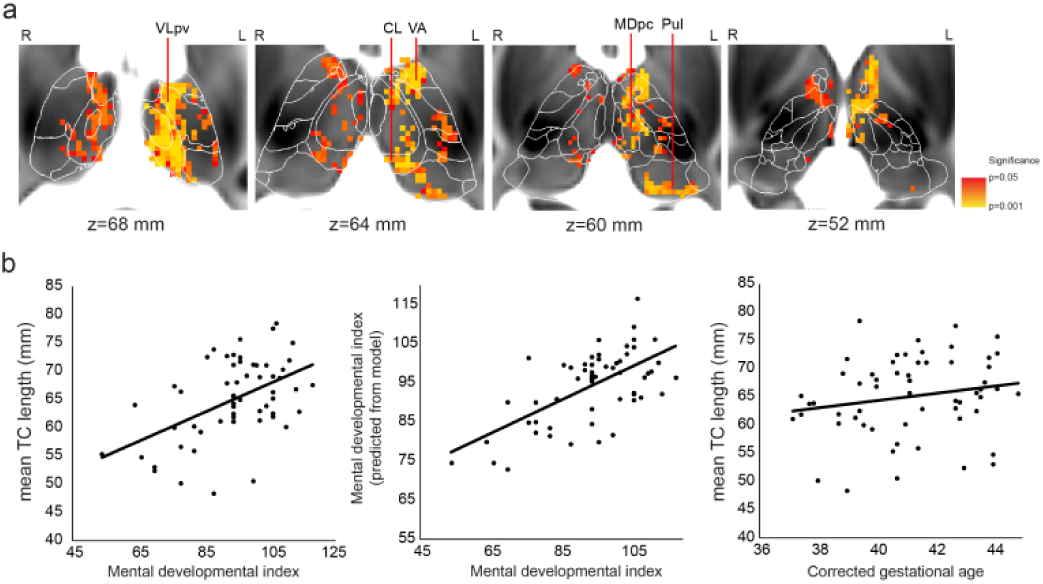
Localization of thalamocortical connections associated with 2-year outcome: thalamic specificity of our findings. (a) Thalamic voxels where TC length was significantly associated with 2-year outcome are displayed as red and yellow overlay (less to more significant) on a T2-weighted MRI template of the 42.week newborn brain, (b) regression plots of the TC length–MDI correlation and gestational age dependence of TC length averaged over the significant voxels. Thalamic nuclei abbreviations are given in **Supplementary Table 2**.

The effect was mostly localized to the ventral-anterior and ventrolateral thalamic nuclei, with smaller clusters found in the centro-lateral (CL) and pulvinar nuclei. Interestingly, more regions were correlated with MDI in the left thalamus than in the right, and the significance levels were also higher (Figure 3/a). The psychomotor development index (PDI) was not correlated with TC length in a model adjusted for PMA. Further analysis of the clinical and demographic variables found a similar relationship between the TC length and MDI: SES, PMA at scanning, and the total time of mechanical ventilation were entered into the model in stepwise linear regression (Table 6, results for all variables including the ones not entered into the model: **Supplementary Table 3)**. The findings on the intrathalamic distribution of voxels in which the TC length was correlated with MDI are summarized in **Table 7**.

**Table 6.** Correlation between mean thalamocortical fiber length (TC length) and mental development index at 2 years after adjusting for clinical and demographic variables.

| Predictors† | Standardized Beta | T | p | Zero-order correlation | Partial correlation | Part correlation |
| --- | --- | --- | --- | --- | --- | --- |
| Mean TC length‡ | 0.460 | 4.481 | .000 | 0.508 | 0.543 | 0.440 |
| Time of mechanical ventilation | -0.354 | -3.529 | .001 | -0.402 | -0.454 | -0.347 |
| SES | -0.308 | -3.023 | .004 | -0.463 | -0.400 | -0.297 |
| PMA | -0.223 | -2.212 | .032 | -0.082 | -0.304 | -0.217 |

† Overall model fit: $F = 17.011$ , adjusted $R^2 = 0.552$ , $P < 0.001$ .
‡ Mean TC calculated from the TFCE-adjusted significant clusters ( $p < 0.05$ )

**Table 7.**
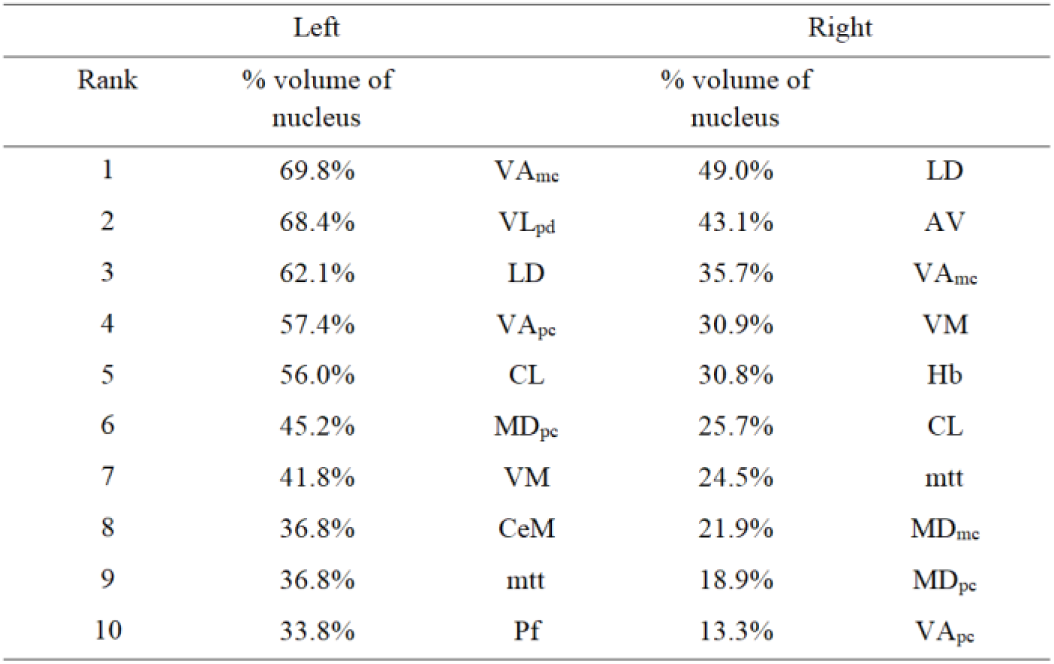
Intrathalamic specificity of our findings: TC length significantly correlated with MDI. Abbreviations of the thalamic nuclei are given in **Supplementary Table 2**.

## Discussion

Our analysis revealed strong correlations between cognitive performance at 2 years, weaker TC strength, and shorter TC length in infants born preterm. Our findings imply that injury or dysmaturation of thalamocortical fiber pathways that may emerge from ventral and nonspecific parts of the thalamus are associated with mental development (MDI), but not psychomotor development (PDI). This provides further evidence for alterations in thalamocortical connectivity after preterm birth [Ball et al., 2013; Fischi-Gomez et al., 2016a; Kelly et al., 2016; Pandit et al., 2014; van den Heuvel et al., 2015]. Our work is an independent validation of a previous study describing a correlation between TC strength and 2-year mental development [Ball et al., 2015]. Remarkably, our findings may indicate a stronger predilection for such association with the frontal and parietal cortical regions. The cortical regions that correlated with cognition at two years of age were more confined to the frontal cortex, and the TC length was significantly associated with MDI in the ventrolateral, ventral anterior, laterodorsal and numerous nonspecific thalamic nuclei.

This relationship appears to be affected by parents’ SES and duration of mechanical ventilation during the NICU stay. SES has previously been confirmed to be an important mediator of cognitive and language development in newborns [Benavente-Fernandez et al., 2019]. Duration of mechanical ventilation has been associated with more brain abnormalities [Brouwer et al., 2017], with white matter abnormalities [Anjari et al., 2009; Ball et al., 2010] and more recently, with alterations in fiber density [Pecheva et al., 2019]. Such comorbidities contribute to impaired WM development and, as confirmed by our post hoc statistical analysis, might be regarded as partial mediators of the main effect found in our analysis. The duration of mechanical ventilation could be also a surrogate marker of the overall level of sickness and cardiorespiratory instability of the study subjects during neonatal course, which also may affect brain integrity.

The thalamus actively participates in cognitive processes by relaying pathways to association cortices, and hence injury to nonspecific or association thalamic nuclei impacts cognitive capabilities. This is reflected by the observation that prematurity affects the thalamocortical connectome [Ball et al., 2013] and cortical microstructure [Ball et al., 2012; Ball, Srinivasan et al., 2013], and the degree of brain injury predicts later cognitive development [Berman et al., 2005; Gui et al., 2018; Little et al., 2010; L. J. Woodward et al., 2006; L. J. Woodward et al., 2011].There is evidence for a close relationship between thalamus injury and impaired cognition in prematurity [Ball et al., 2015]. Lower cognitive performance at 2 years of age was not only found to be linked not only to a global reduction of thalamocortical connectivity [Ball et al., 2015] but also to lower neuronal metabolite concentrations in the thalamus [Hyodo et al., 2018]. Reduced thalamus volume is associated with decreased structural integrity of posterior fiber pathways in the brain of school-age children born prematurely. In severe cases with periventricular leukomalacia, thalamic volume reduction was found to have an effect on general intelligence and working memory measures [Zubiaurre-Elorza et al., 2012]. The thalamus is an important relay station to cortical regions involved in higher cognition [Parnaudeau et al., 2018]. This is confirmed by studies that found association between the circuitry of the mediodorsal nucleus (MD) and executive functions [Ardila, 2019; Jakab et al., 2012].

While there is no strong support for premature birth selectively affecting neuronal circuits, our findings may shed light on a way in which cognitive development becomes disrupted due to increased vulnerability of its neural correlates to injury after premature birth. Our study population included infants born between the 26^th^ and 31^st^ weeks of gestation. During this period, the exposure of the immature brain to various toxic events in the neonatal intensive care unit and critical developmental processes such as myelination and the emergence of thalamocortical connections occur. By this time, the first TC fibers have already reached the cortex [Kostovic and M. Judas, 2002; Kostovic and N. Jovanov-Milosevic, 2006], and the brain has progressed to a stage of rapid development, characterized by nonlinear increase in brain volume and an increase in synaptic density. Human thalamocortical axons show prolonged growth (4 months), and somatosensory fibers precede the ingrowth of fibers destined for frontal and occipital areas [Krsnik et al., 2017]. At term-equivalent age, the topography of thalamocortical connections largely resembles that of an adult [Ferradal et al., 2019]. Thus, there is overwhelming neuroanatomical evidence for the rapid and late development of the frontal lobe in the human brain [Hodel, 2018], which confers increased vulnerability to adverse environments. Fetal studies have characterized a posterior-to-frontal (sensory-to-higher association) gradient in functional brain development [Jakab et al., 2014; Thomason et al., 2013], meaning that regional differences in TC development between the 27^th^ and 31^st^ weeks of gestation may open a window of selective vulnerability of frontal and parietal circuitry. During the early postnatal period in preterm infants, frontal lobe myelination differs from that in other brain regions by the longer persistence of premyelinating oligodendrocytes [Back et al., 2001]; these are more vulnerable than mature oligodendrocytes to perinatal insults [Liu et al., 2013]. This observation allows us to speculate that impaired frontal lobe myelination, presumably also affecting the anterior thalamic radiation, would result in lower structural connectivity indices, as tractography becomes more uncertain at lower anisotropy values.

The following limitations of our study merit mentioning. The generalizability of our results is limited by the relatively low case number (n=58), which reduces the statistical power of multivariate linear regression analysis with multiple covariates in the model. Furthermore, long-term follow-up results are necessary to confirm whether the association revealed between TC, TC length, and mental developmental index is sustained in later stages of development. Probabilistic thalamocortical tractography faces inherent technical limitations. First, afferent and efferent thalamic pathways are indistinguishable because diffusion direction does not respect tract polarity. Probabilistic tractography becomes increasingly uncertain at the WM/gray matter interface and in regions with lower diffusion anisotropy, such as the thalamus. While numerous reports have confirmed the value of this method for describing the connectional topography of the human thalamus [Behrens et al., 2003; Ferradal et al., 2019], it is likely that the current spatial and angular resolution is insufficient to identify some of the nonspecific nuclei of the thalamus, as fibers emerging from these must cross adjacent specific thalamic nuclei encompassing highly ordered, white-matter-rich regions. Despite these limitations, we rely confidently on the ability of probabilistic tractography for mapping thalamus-frontal and prefrontal connectivity based on previous neuroanatomical and imaging studies [Jakab et al., 2012; Klein et al., 2010; Le Reste et al., 2016].

The use of TC length as indicator of thalamocortical connectivity strength is not straightforward because it may not only reflect physical distance. First, TC length was measured in standard anatomical space, which reduces the effect of individual size variability. Shorter length may be caused by the shorter distance that probabilistic tracing particles on average traverse if the connection probability is lower, and by the same token, pathways are terminated earlier if FA is lower. TC and TC length should therefore be interpreted as variables that do not directly reflect the number of mono- or polysynaptic pathways reaching the cortex, which would be the simplest definition of anatomical connectivity strength, but a variety of diverse effects. These include the cumulative effect of diffusion anisotropy and fiber uncertainty along the pathways, which in turn may also reflect myelination and the actual number of axons in a voxel.

The complete lack of histologically defined, human newborn thalamus atlases increases the difficulty of interpreting our results on thalamic specificity. While structural and functional connectivity at birth appears to largely overlap with adult patterns, the precise variability of this in preterm infants remains to be characterized. Our previous report using the shape-model-based alignment of the thalamus atlas was found to be flexible enough to tackle large shape variability, as proven by matching the atlas to high-resolution MRI of postmortem, fixed thalamus samples [Jakab et al., 2012].

## Conclusions

Our study confirmed the value of TC circuitry in predicting cognitive development in preterm born infants. Cognitive abilities are important for school and academic success and ultimately for quality of life. Hence, early identification of aberrant brain development that underlies later cognitive deficits is essential to start early interventions and to monitor the effects and efficacy of intervention on brain development.

## Acknowledgements

A.J. was supported by the OPO Foundation, the Forschungszentrum für das Kind (FZK) Grant, the Foundation for Research in Science and the Humanities at the University of Zurich, the Anna Müller Grocholski Foundation, the EMDO Foundation and the Prof. Dr. Max Cloetta Foundation. R.T. was supported by the EMDO Foundation. The study was supported by a grant received from the Swiss National Science Foundation (3200B0-108176) and was registered at clinicaltrials.gov (identifier: NCT 00313946).

We would like to thank Simon Milligan for his help in language editing the manuscript. The authors declare that they do not have conflict of interest.

**This manuscript is the author’s original version**.

## Supplementary Tables and Figures

**Supplementary Figure 1.**
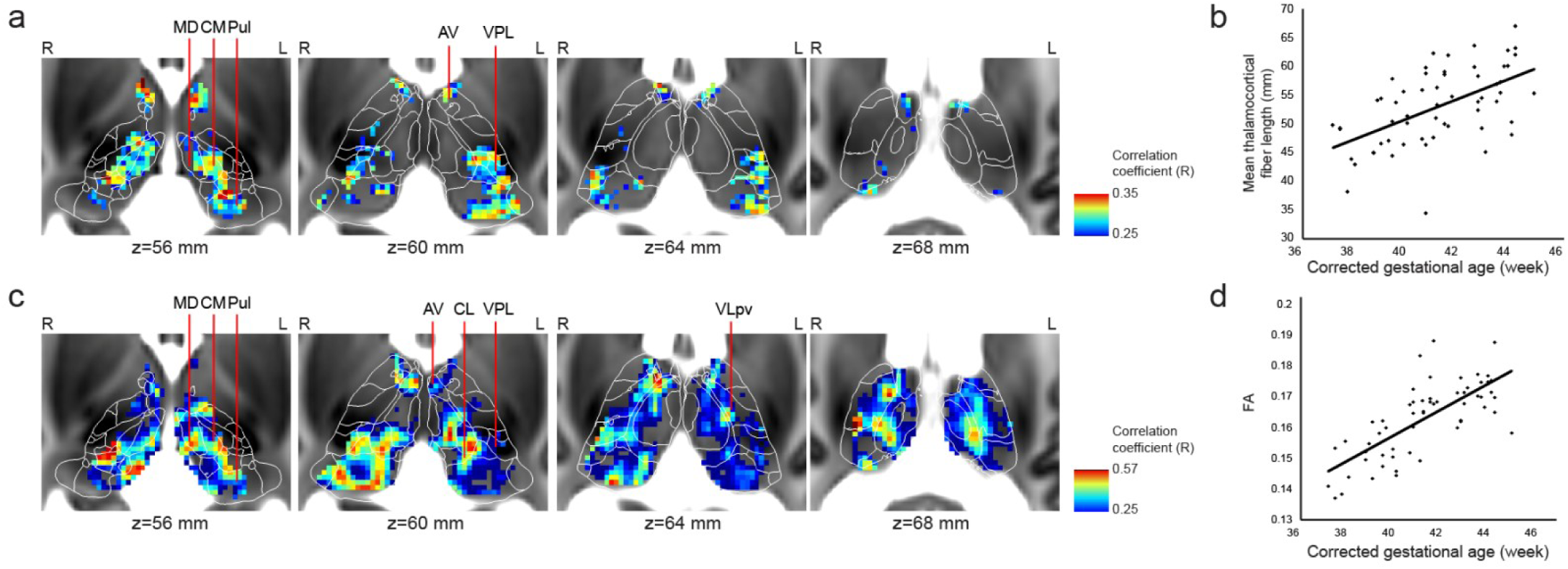
Gestational age-dependent change of fractional anisotropy and TC length. A multivariate linear regression analysis was performed for each voxel in the thalamus to reveal the association between corrected gestational age and FA and TC length.

**Supplementary Table 1.** Post hoc stepwise linear regression analysis of the relationship between TC and MDI: all clinical and demographic variables.

| Variable | Beta | t | p | Partial correlation | Entered into model |
| --- | --- | --- | --- | --- | --- |
| GA at birth | .220 | 1.859 | .069 | .257 | No |
| PMA at scanning | -.210 | -1.724 | .091 | -.239 | Yes |
| Birth weight | .200 | 1.623 | .111 | .226 | No |
| Birth weight, Z-Score | .151 | 1.218 | .229 | .171 | No |
| Head circumference | .169 | 1.379 | .174 | .193 | No |
| Head circumference, Z-Score | .131 | 1.054 | .297 | .149 | No |
| Parental SES | -.375 | -3.327 | .002 | -.429 | Yes |
| BPD grade | -.320 | -2.819 | .007 | -.374 | No |
| Duration of mechanical ventilation | -.405 | -3.760 | .000 | -.473 | Yes |
| Sepsis (Y/N) | -.278 | -2.378 | .021 | -.322 | No |
| EPO treatment group | .175 | 1.433 | .158 | .201 | No |
| Sex | -.061 | -.496 | .622 | -.071 | No |

**Supplementary Table 2.**
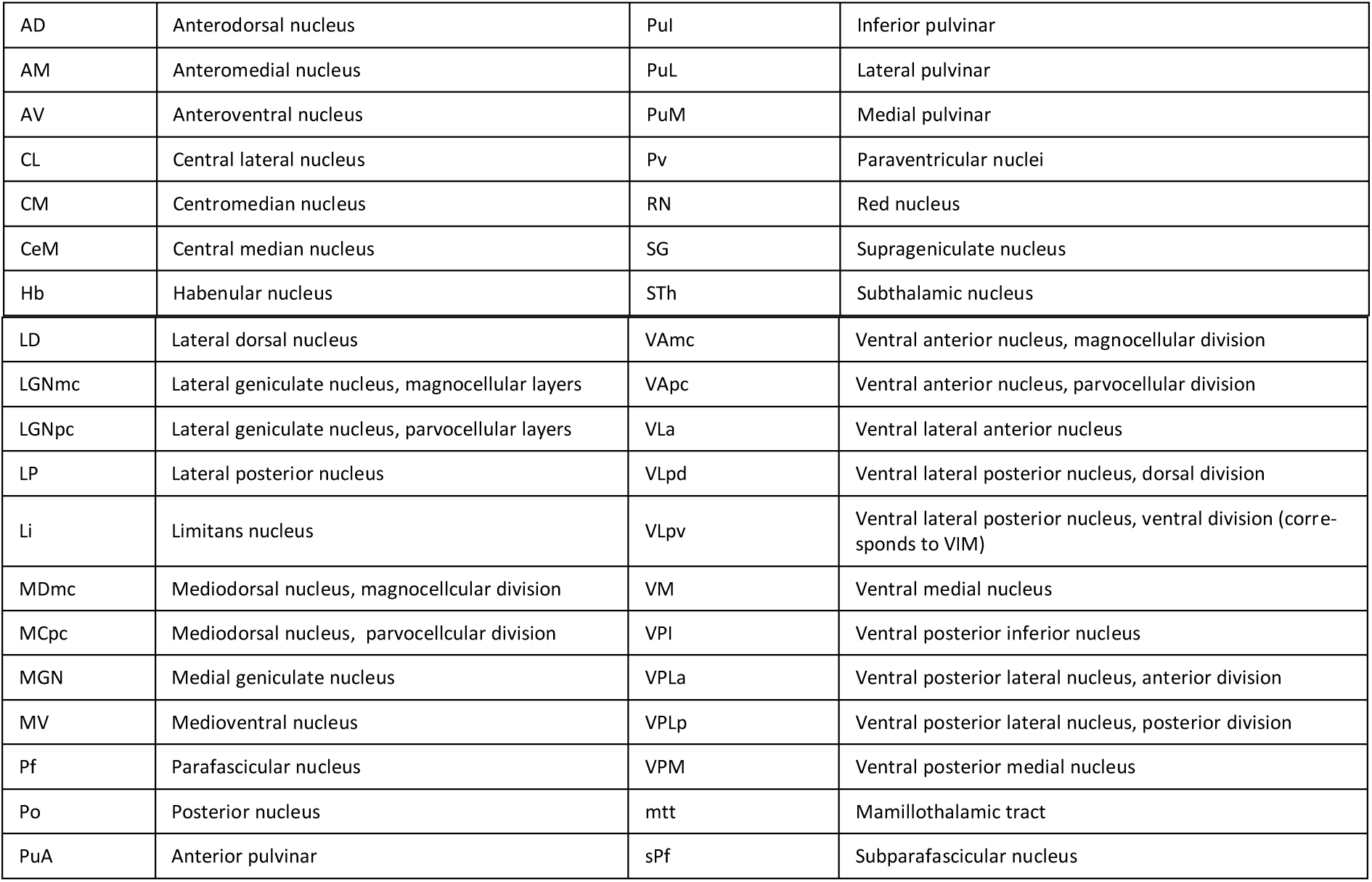
Thalamic nuclei abbreviations.

**Supplementary Table 3.** Post hoc stepwise linear regression analysis of the relationship between TC length and MDI: all clinical and demographic variables.

| Variable | Beta | t | p | Partial correlation | Entered into model |
| --- | --- | --- | --- | --- | --- |
| GA at birth | .163 | 1.347 | .184 | .189 | No |
| PMA at scanning | -.184 | -1.504 | .139 | -.210 | Yes |
| Birth weight | .058 | .471 | .640 | .067 | No |
| Birth weight, Z-Score | .039 | .319 | .751 | .045 | No |
| Head circumference | .052 | .421 | .676 | .060 | No |
| Head circumference, Z-Score | .013 | .106 | .916 | .015 | No |
| Parental SES | -.367 | -3.198 | .002 | -.416 | Yes |
| BPD grade | -.267 | -2.234 | .030 | -.304 | No |
| Duration of mechanical ventilation | -.373 | -3.357 | .002 | -.432 | Yes |
| Sepsis (Y/N) | -.246 | -2.044 | .046 | -.280 | No |
| EPO treatment group | .188 | 1.546 | .129 | .216 | No |
| Sex | -.036 | -.294 | .770 | -.042 | No |

